# A three-country analysis of the gut microbiome indicates taxon associations with diet vary by location and strain

**DOI:** 10.1101/2024.10.01.616090

**Authors:** Lora Khatib, Se Jin Song, Amanda H Dilmore, Jon G Sanders, Caitriona Brennan, Alejandra Rios Hernandez, Tyler Myers, Renee Oles, Sawyer Farmer, Charles Cowart, Amanda Birmingham, Edgar A. Diaz, Oliver Nizet, Kat Gilbert, Nicole Litwin, Promi Das, Brent Nowinski, Mackenzie Bryant, Caitlin Tribelhorn, Karenina Sanders-Bodai, Soline Chaumont, Jan Knol, Guus Roeselers, Manolo Laiola, Sudarshan A. Shetty, Patrick Veiga, Julien Tap, Muriel Derrien, Hana Koutnikova, Aurélie Cotillard, Christophe Lay, Armando R. Tovar, Nimbe Torres, Liliana Arteaga, Antonio González, Daniel McDonald, Andrew Bartko, Rob Knight

## Abstract

Emerging research suggests that diet plays a vital role in shaping the composition and function of the gut microbiota. While significant efforts have been made to identify general patterns linking diet to the gut microbiome, much of this research lacks representation from low- and middle-income countries such as Mexico. Additionally, both diet and the gut microbiome have highly complex and individualized configurations, and there is growing evidence that tailoring diets to individual gut microbiota profiles may optimize the path toward improving or maintaining health and preventing disease. Using fecal metagenomic data from 1,291 individuals across three countries, we examine two bacterial genera prevalent in the human gut, *Prevotella* and *Faecalibacterium*, which have gained significant attention due to their potential roles in human health. We find that they show significant associations with many aspects of diet, but that these associations vary in scale and direction, depending on the level of metagenomic resolution and the contextual population. These results highlight the growing importance of assembling metagenomic datasets that are standardized, comprehensive, and representative of diverse populations to increase our ability to tease apart the complex relationship between diet and the microbiome.

## Importance

An analysis of fecal microbiome data from individuals in the United States (US), United Kingdom (UK), and Mexico shows that associations with dietary components vary both by country and by level of resolution. Our work sheds light on why there may be conflicting reports regarding microbial associations with diet, disease, and health.

## Observation

In this study, we explored the relationships between the gut microbiome at different levels (taxonomic species and metagenome-assembled genomes (MAGs)) and various dietary factors, including dietary patterns, nutrients, and food groups, using data collected from subjects in three countries (Figure 1a-b). We conducted metagenomic sequencing of fecal samples from 442 adult participants in the US, 342 in the UK, and 507 in Mexico, recruited to participate through the Microsetta Initiative platform(1). We evaluated long-term dietary intake using the VioScreen™ food frequency questionnaire (FFQ) (Version 5, VioCare, Princeton, NJ)(2). We adapted this version for Mexico to include common foods in the region (Version 5-Mex).

**Figure 1.** Study and data overview. a. The table shows the number of samples analyzed by shotgun metagenomics from each country and the resulting sample size after filtering steps. b. Data collected included taxonomic profiles and metagenome-assembled genomes from the sequence data, and diet and lifestyle information from self-reported answers to questionnaires. c. Radar plots show z-normalized values for key personal and diet-related variables, averaged by country. d. A principal coordinates analysis plot shows robust PCA distance among the microbiome samples using the taxonomic data, colored by the cohort country. Partial R^2^ from a PERMANOVA is reported. A confusion matrix shows the classification accuracy using a random forest classifier (5-fold cross validation) on the taxonomic feature table.

Consumption of food groups expressed in Kcal/day was derived from raw VioScreen™ entries and several filtering criteria were applied to ensure data quality based on the number of items consumed, total energy intake and the time of FFQ completion as previously described(3).

All cohorts consisted of more females than males with an average BMI of 25(± 5 SD), but varied in most other characteristics, with subjects in the UK being generally older in age, and subjects in Mexico reporting higher incidence of diabetes and IBD, as well as having the lowest diet quality (based on the Healthy Eating Index, HEI)(Figure 1c, Table 1). However, all three cohorts constituted individuals healthier than the general US population, reporting a lower BMI, higher HEI, and lower incidence of diabetes and cardiovascular disease(4, 5).

**Table 1.** Key demographic, health, and diet characteristics of the three cohorts.

|  | Mexico | UK | US | statistic | p_value |
| --- | --- | --- | --- | --- | --- |
| No. of participants | 460 (39.1%) | 307 (26.1%) | 410 (34.8%) | 31.026 | 1.83E-07 |
| Age | 42.195 ± 15.133 | 52.116 ± 13.269 | 45.011 ± 15.22 | 81.618 | 1.89E-18 |
| Sex: Female | 307 (66.7%) | 223 (72.6%) | 262 (63.9%) | 6.191 | 0.04525496 |
| Education level: College graduate | 206 (45.8%) | 153 (50.2%) | 189 (46.8%) | 20.238 | 0.00044809 |
| Alcohol Frequency: Regularly | 36 (7.9%) | 80 (26.4%) | 92 (22.4%) | 52.158 | 4.72E-12 |
| BMI | 25.573 ± 5.01 | 25.091 ± 5.706 | 25.23 ± 5.314 | 5.114 | 0.0775445 |
| Diabetes | 31 (6.9%) | 9 (3.0%) | 15 (3.7%) | 7.777 | 0.02047483 |
| CVD | 9 (2.0%) | 14 (4.6%) | 16 (3.9%) | 9.818 | 0.04360907 |
| Autoimmune Disease | 19 (4.3%) | 51 (17.0%) | 52 (12.8%) | 34.029 | 4.08E-08 |
| IBD | 64 (15.1%) | 9 (3.0%) | 9 (2.3%) | 61.016 | 5.63E-14 |
| IBS | 123 (28.4%) | 82 (27.1%) | 72 (18.3%) | 12.9 | 0.00158089 |
| Gluten intolerance | 26 (6.3%) | 31 (10.3%) | 51 (12.9%) | 23.303 | 0.00070106 |
| Lactose intolerance | 136 (31.1%) | 29 (9.9%) | 74 (18.6%) | 49.665 | 1.64E-11 |
| Bowel movement, normal | 349 (77.0%) | 218 (73.4%) | 300 (76.5%) | 30.367 | 3.35E-05 |
| Diet Type (Vegetarian) | 6 (1.3%) | 21 (6.9%) | 27 (6.6%) | 83.655 | 8.96E-15 |
| Plant Diversity (More than 20) | 55 (12.1%) | 105 (34.9%) | 96 (23.5%) | 189.206 | 1.20E-36 |
| Vegetable Frequency, Regularly | 358 (78.0%) | 286 (93.5%) | 345 (84.8%) | 37.794 | 1.24E-07 |
| Fruit Frequency, Regularly | 314 (68.6%) | 207 (67.9%) | 251 (61.5%) | 6.348 | 0.17461969 |
| Whole Grain Frequency, Regularly | 194 (42.7%) | 155 (50.7%) | 243 (59.7%) | 25.219 | 4.55E-05 |
| Red Meat Frequency, Regularly | 171 (37.6%) | 46 (15.0%) | 60 (14.7%) | 90.161 | 1.22E-18 |
| Milk/Cheese Frequency, Regularly | 303 (66.0%) | 188 (61.2%) | 198 (48.9%) | 28.107 | 1.19E-05 |
| Sugar Sweetened Beverage Frequency, Regularly | 46 (10.1%) | 7 (2.3%) | 13 (3.2%) | 76.907 | 7.87E-16 |
| Energy intake from diet(kcal/day) | 2284.797 ± 901.074 | 2007.523 ± 688.092 | 2017.892 ± 678.689 | 21.896 | 1.76E-05 |
| Ultraprocessed Foods (kcal/day) | 345.554 ± 127.168 | 438.919 ± 141.507 | 487.323 ± 152.632 | 195.972 | 2.79E-43 |
| Processed Foods (kcal/day) | 213.998 ± 100.985 | 80.455 ± 62.177 | 77.713 ± 59.797 | 493.984 | 5.40E-108 |
| Minimally processed Foods (kcal/day) | 388.19 ± 121.734 | 411.377 ± 130.741 | 359.726 ± 133.215 | 34.562 | 3.13E-08 |
| Carbohydrate (%) | 47.283 ± 7.625 | 46.105 ± 9.872 | 44.178 ± 9.259 | 28.152 | 7.71E-07 |
| Fat (%) | 34.088 ± 6.295 | 34.988 ± 8.698 | 36.854 ± 8.159 | 30.223 | 2.74E-07 |
| Protein (%) | 16.307 ± 3.189 | 14.753 ± 3.007 | 15.341 ± 3.526 | 47.901 | 3.97E-11 |
| Healthy Eating Index (2010) | 64.741 ± 9.495 | 73.197 ± 10.537 | 72.456 ± 10.516 | 182.008 | 3.00E-40 |
| Healthy Eating Index (2015) | 63.348 ± 9.308 | 71.027 ± 10.242 | 70.166 ± 10.398 | 147.01 | 1.19E-32 |

Stool collection and DNA extraction were performed similarly across the three cohorts. Sample libraries generated an average of 4,559,420 sequences per sample(± 1,750,993 SD), which were then quality controlled and human sequence filtered following the recommendations from Armstrong et al. (2022)(6), before generating operational genomic unit (OGU) through the woltka pipeline(7) (Figure 1b). A robust PCA analysis(8) showed that the strongest effect on microbiome composition is geographic location (PERMANOVA pseudo-F=133.8, *p*=0.001; Figure 1d), such that a random forest classifier could predict location with very high accuracy (mean AUC: 1.00 for Mexico, 0.95 for UK, 0.96 for USA) across 5-fold cross-validation. The next top variables (by PERMANOVA Partial R^2^ value) showing significant effects not related to diet were BMI (R^2^=0.007745, p=0.001), antibiotic history (R^2^=0.004224, p=0.001), and mental and physical well-being (R^2^=0.002114, p=0.034). Therefore, all downstream analyses, where possible, included these factors as covariates.

To assess whether sub-species-level genetic diversity was associated with diet, we constructed contigs followed by binning using the method described in Sanders et al. (2023)(9). From this set, we derived species-level genome bins (sGB) using dRep(10) to ensure distinct species representatives. Next, we calculated nucleotide diversity within each sGB for each individual using InStrain(11), and then down-selected from 286 dietary variables (representing estimated micro and macronutrients) using linear LASSO regression followed by linear mixed effects models. This investigation revealed that *Prevotella* and *Faecalibacterium* nucleotide diversity showed the strongest associations with dietary variables (Figure 2a). Specifically, we found that *Prevotella* nucleotide diversity is positively associated with the consumption of whole grains (*t*=4.962, p=7.21e-7) and negatively associated with time since antibiotic use (*t*=-6.949, p=4.17e-12). The nucleotide diversity of *Faecalibacterium* showed a positive association with dietary fiber (*t*=9.282, p=2.20e-20), consistent with previous studies(12), and, unexpectedly, with processed meat (*t*=9.060, p=1.67e-19). We suspect this association is related to participants in Mexico who self-reported high fiber intake along with high amounts of processed food, including processed meat.

**Figure 2.** Aspects of *Prevotella* and *Faecalibacterium* at different levels of resolution highlight the complexity of diet-microbe associations, each showing ties with many dietary variables, but varying in direction and strength by country. a. Heatmap shows t-scores and p-values for testing the correlation between the nucleotide diversity in species-level Genome Bins (sGB)) and dietary variables, pooled per individual by bacterial genus. b. Scatter plots show the dietary variables with the strongest negative and positive relationships with *Prevotella* (top two panels) and *Faecalibacterium* (bottom two panels). Points are colored by cohort country, with corresponding correlation lines. Correlation coefficients and p-values are shown in the insets. c. An association map shows the dietary variables that were significantly correlated with the log ratio of each *Prevotella* sGB to the sum of all *Prevotella* (top) and each *Faecalibacterium* sGB to the sum of all *Faecalibacterium* (bottom) in the full dataset and stratified by country, after accounting for the variation explained by covariates (cohort, BMI, antibiotic history, level of well-being). Multiple comparisons were corrected using the Benjamin-Hochberg method with a 5% FDR.

We then investigated whether the log ratio of these genera against *Bacteroides*, the most prevalent taxon, showed a similar pattern to nucleotide diversity. We found that when collapsed on the genus level, we detected significant associations with many of the expected dietary variables consistent with previous studies, such as an enrichment of both taxa with higher dietary fiber intake (Figure S1). However, the strength of these associations was inconsistent across the three countries, with the strongest associations typically in the US (Figure 2b).

Next, we performed the same analysis on the sGBs for these two genera, and on a subset of the numeric dietary variables that broadly summarized diet (Table S1). Only 14 out of 56 *Prevotella* sGBs showed significant associations with any of the dietary variables (Figure 2c). Few *Prevotella* sGBs showed consistency across the cohorts. In the full dataset, *P. disiens* was positively associated with alcohol consumption, *P. sp002481295* was associated with animal and protein intake, and two sGBs without named genomes in the Genome Taxonomy Database (GTDB) (*P. MAG sGB 00348* and *P. copri sGB 00022*) were associated with starch intake. However, the remaining 10 sGBs showed associations that were country-specific. For example, *P. timonensis* showed positive associations with vegetable and fiber intake in the US, but not in the UK or Mexico, where it instead showed an association with saturated fat intake.

Finally, and in contrast, nearly all (17 out of 19) of the *Faecalibacterium* sGBs showed significant associations. Several *Faecalibacterium* sGBs (most notably *F. MAG_sGB_00436, F. prausnitzii_I*, and *F. sp900539885*) were positively associated with intake of vegetable, fruit, carbohydrates (including fibers, pectin, and starch), and unprocessed or minimally processed foods, as well as an overall healthy diet (high HEI), while negatively associated with animal protein and dairy. However, other sGBs showed opposite trends (*F. prausnitzii_C, F. MAG_sGB_00406*), associating with animal protein, saturated fat, and an overall poorer diet (low HEI), but only in the UK cohort. Interestingly, no sGBs were found to be significantly associated with any dietary variables in Mexico.

Understanding the complex associations between diet and the microbiome can contribute to our knowledge of gut microbiota modulation and its implications for personalized nutrition. Here, we focused on two bacterial genera, *Prevotella* and *Faecalibacterium*, which have shown the most association with diet and human health across studies. Both taxa can ferment dietary fibers(12-14), and are associated with plant-based foods(15, 16). *Prevotella* has been associated with both beneficial and detrimental health effects. For example, while *Prevotella* is associated with high-fiber diets(14), it has also been associated with inflammatory conditions, such as periodontal disease, rheumatoid arthritis, and metabolic syndrome(17-19). *Faecalibacterium*, by contrast, has consistently been associated with beneficial health effects(20). The strain-level contributions to these associations are not yet fully understood, but growing studies have revealed that *Prevotella* and *Faecalibacterium* are richer in strain diversity than previously appreciated(21, 22). A recent study showed that *Faecalibacterium* strain diversity can vary among people of different ages, populations, lifestyles, and disease status(23), with non-Western populations showing higher prevalence and diversity. In our study, despite more individuals from Mexico having a *Prevotella*-dominant microbiome as well as an enrichment in both *Prevotella* and *Faecalibacterium* compared to the US and UK (Figure S2a-b), they had the lowest diversity of *Prevotella* of the three cohorts (Figure S2c), as well as the lowest intake of fiber and plant-based foods (Table 1). However, they had a higher diversity of *Faecalibacterium* than in the UK cohort. These results suggest that strain diversity in both taxa may be associated with a diet rich in fiber due to increased potential in the metabolism of those substrates, but that individual strains may associate or respond to the same diet in different ways.

Capturing a broader range of human and dietary diversity will help us better untangle the complex relationships between microbial strains and diet. Our study indicates that while some associations with diet are consistent between countries at taxonomic levels, some specific associations are detected only at the strain level. Efforts to develop a better understanding of whether other components of diet or the microbiome modify the relationships between a given strain and a given dietary item in a region-specific manner are warranted.

---

|  | USA | UK | Mexico |
| --- | --- | --- | --- |
| Adult participants | 442 | 342 | 507 |
| <1,000,000 reads | 2 | 2 | 39 |
| no answer to covariate(s)* | 30 | 33 | 8 |
| final n | 410 | 307 | 460 |

\*bmi, antibiotic history, level of well-being

| fecal metagenomic data |  | metadata |  |
| --- | --- | --- | --- |
| taxonomic | assembled | Microsetta<br>questionnaire | Vioscreen food<br>frequency questionnaire |
| 956 OGUs | 16,652 MAGs | lifestyle, diet, personal | nutrients, food items,<br>food groups, dietary<br>patterns |

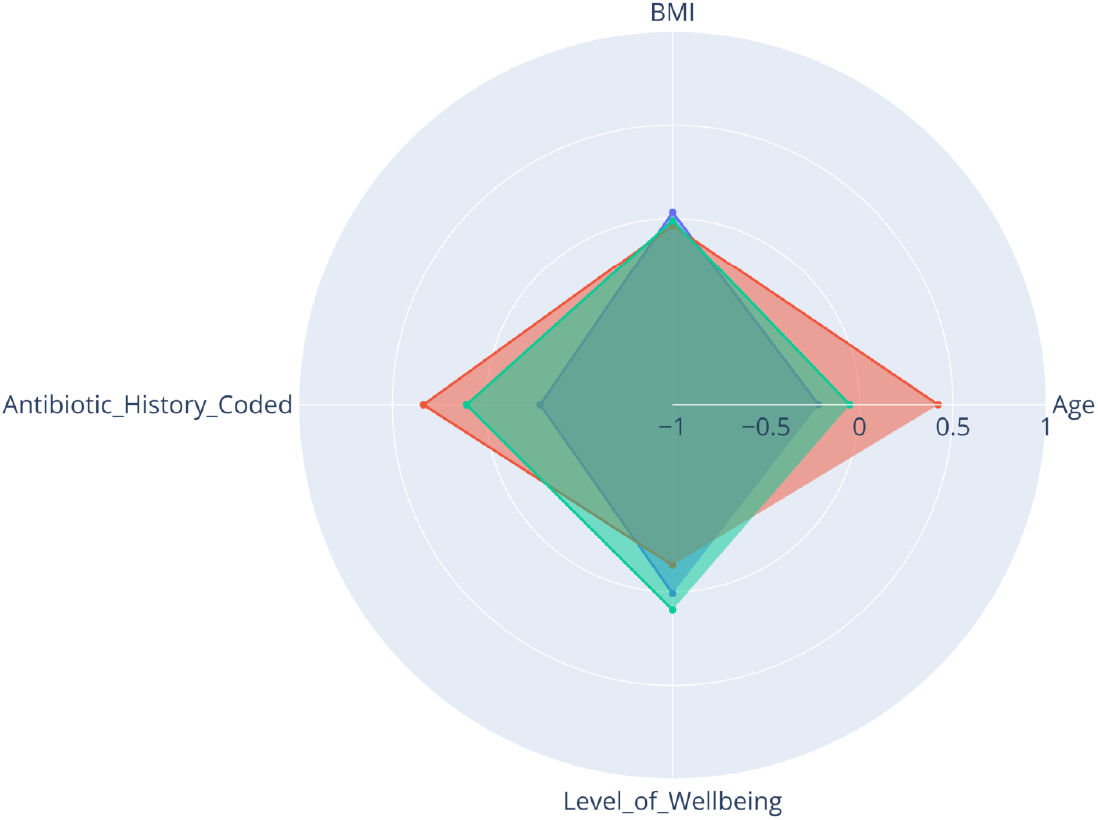
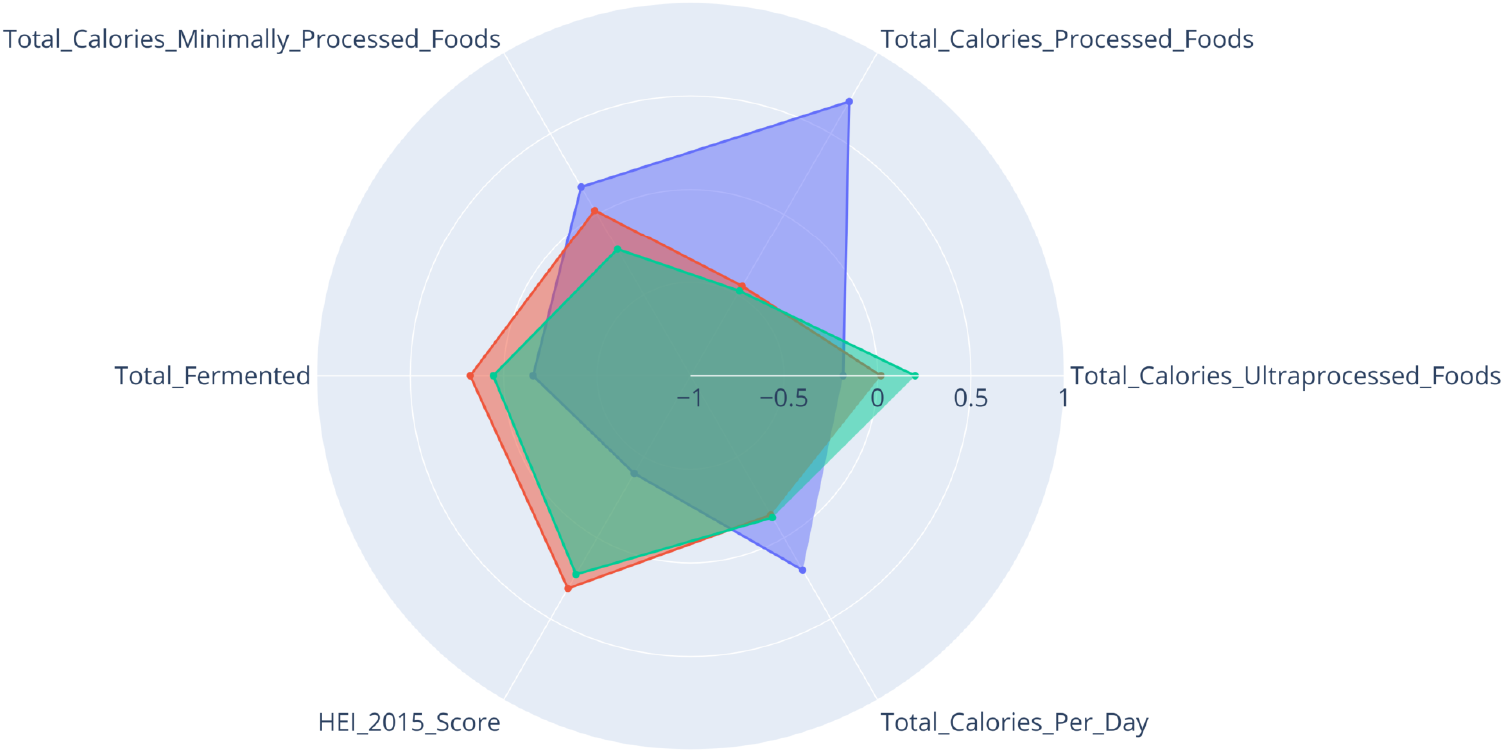
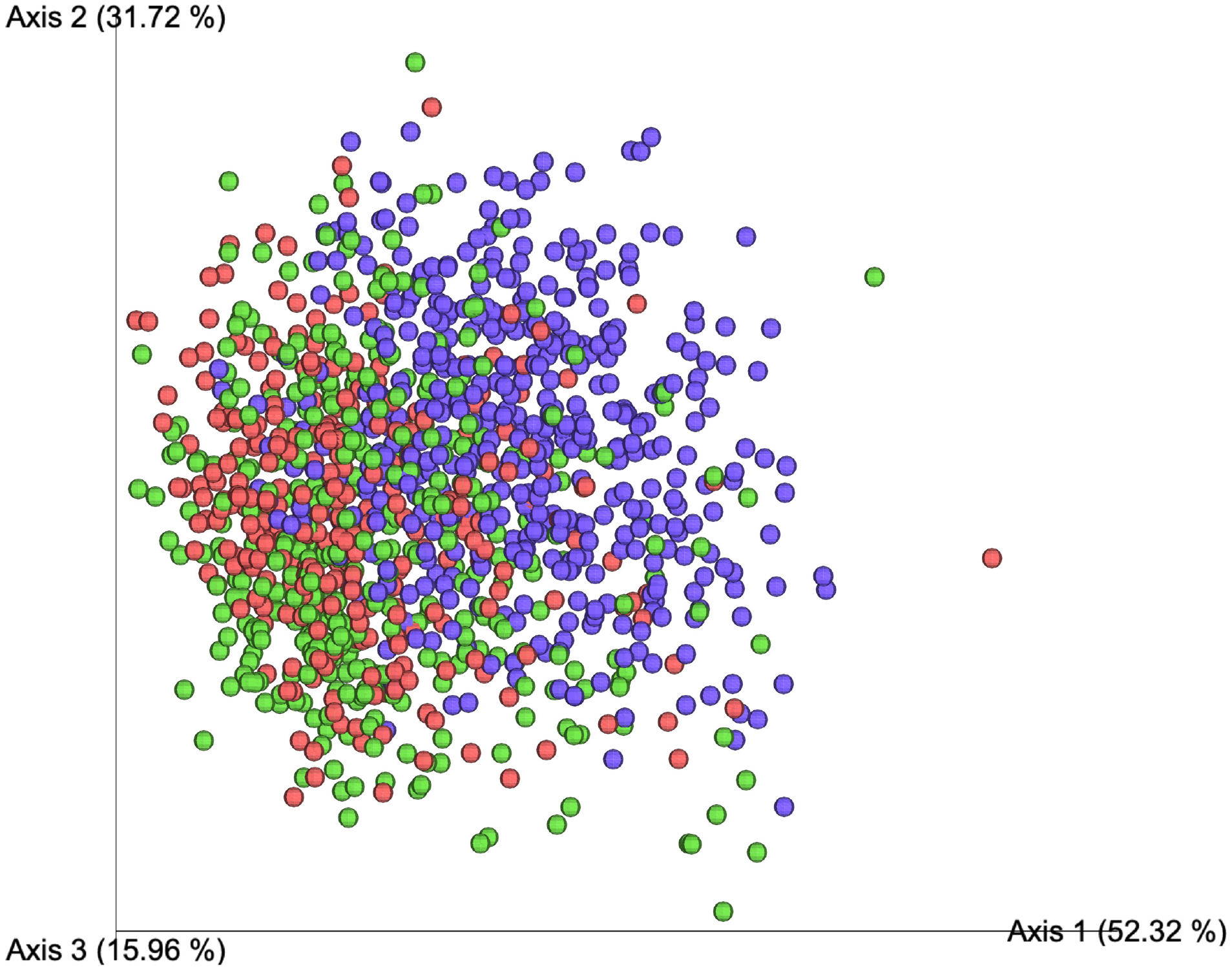
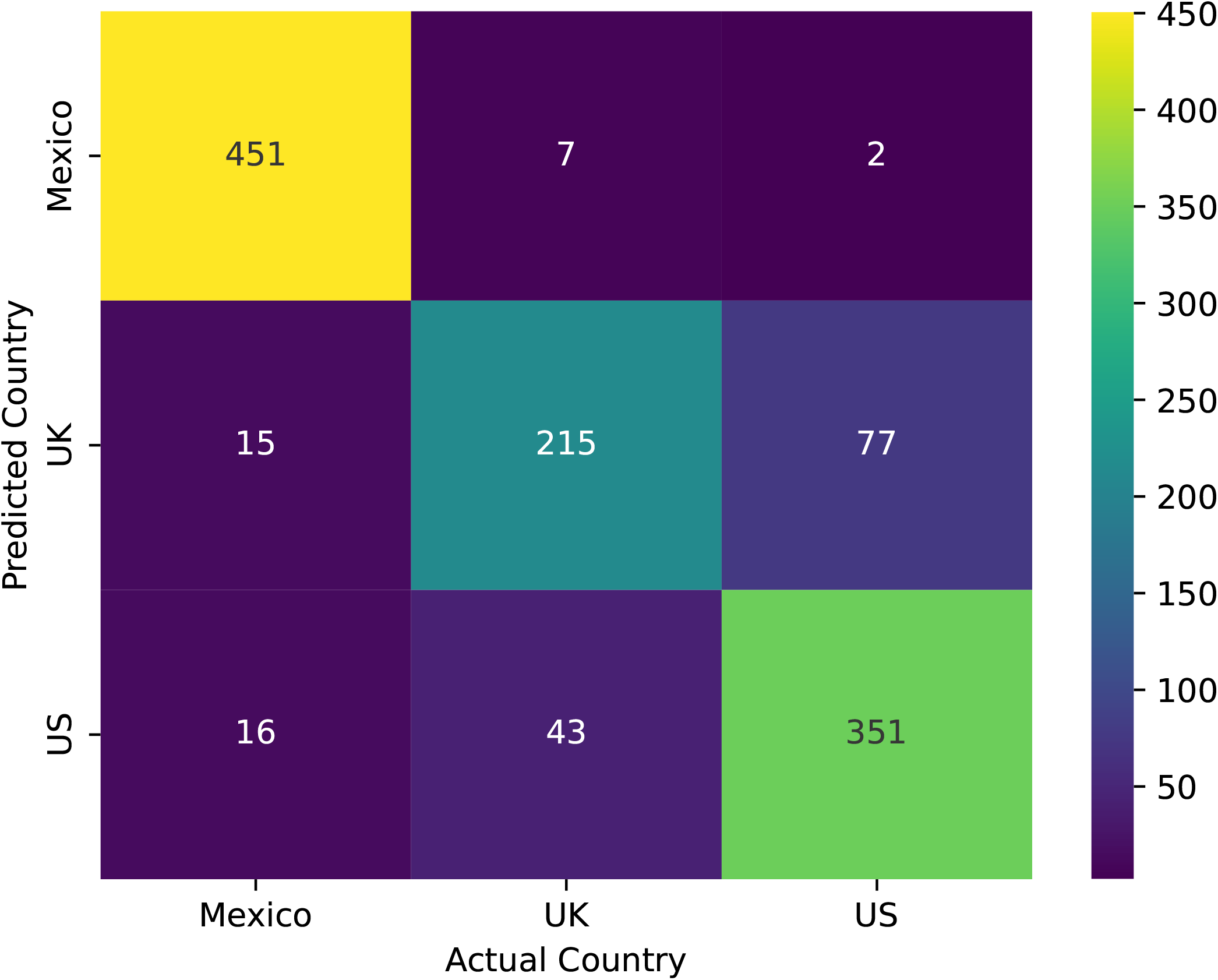

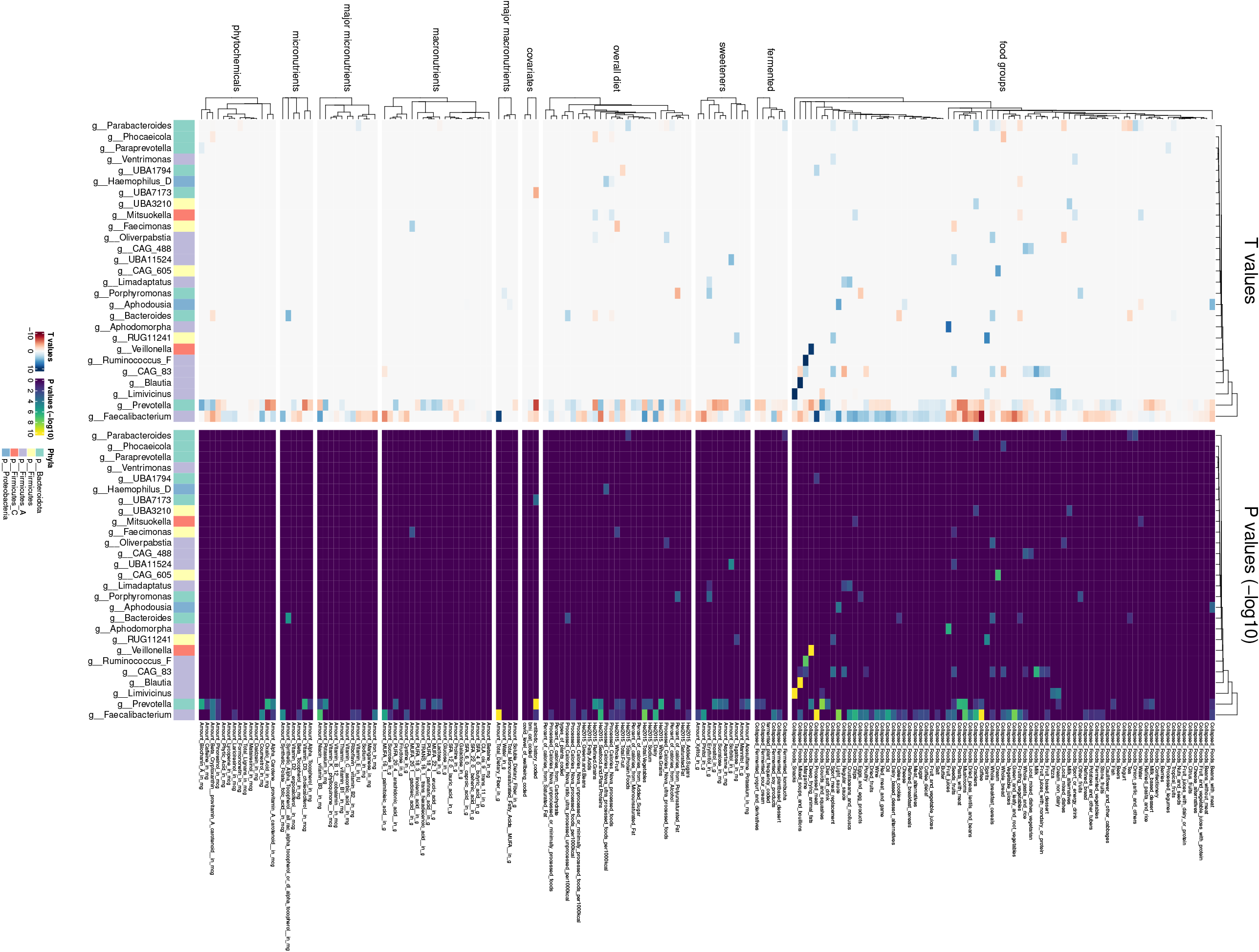
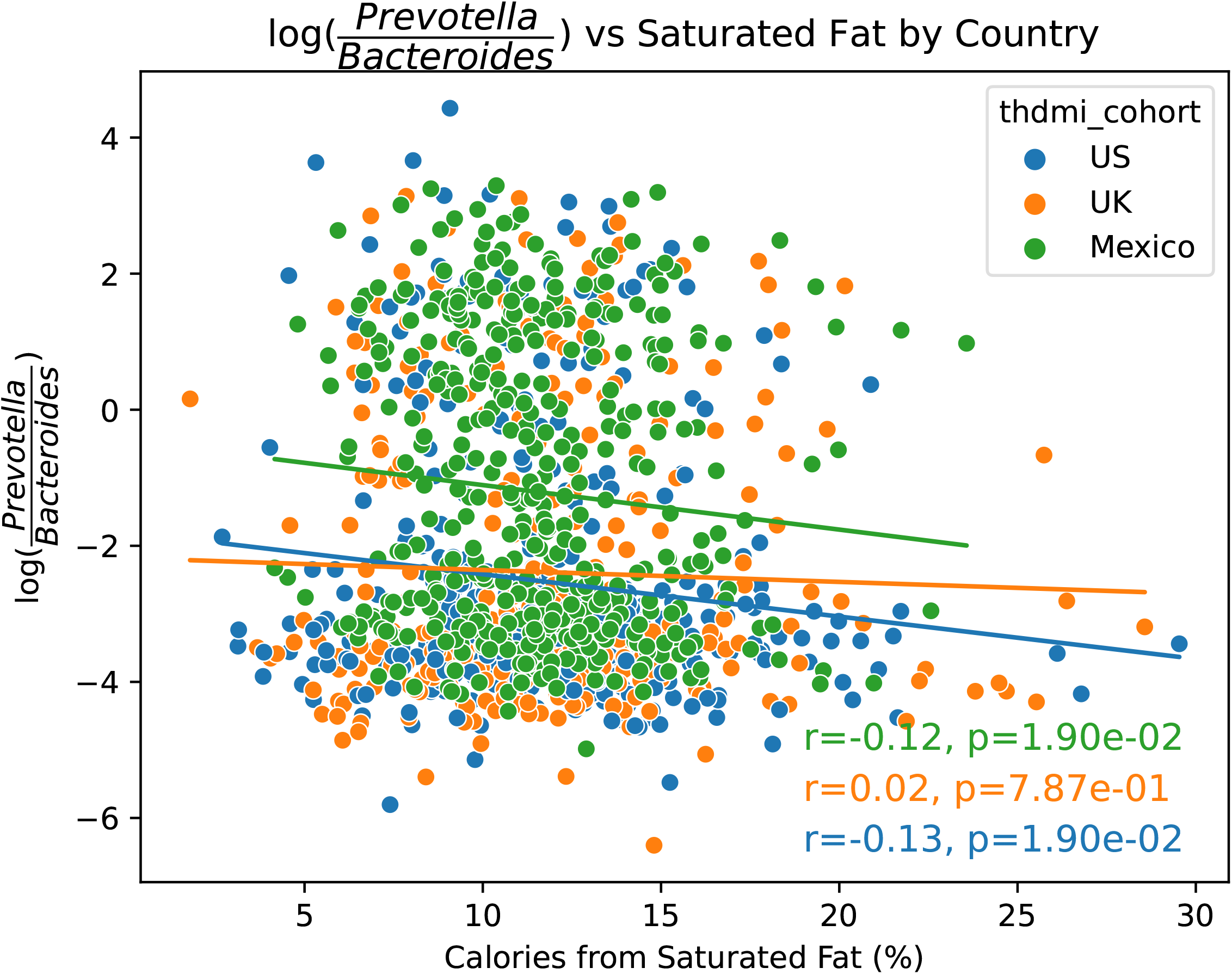
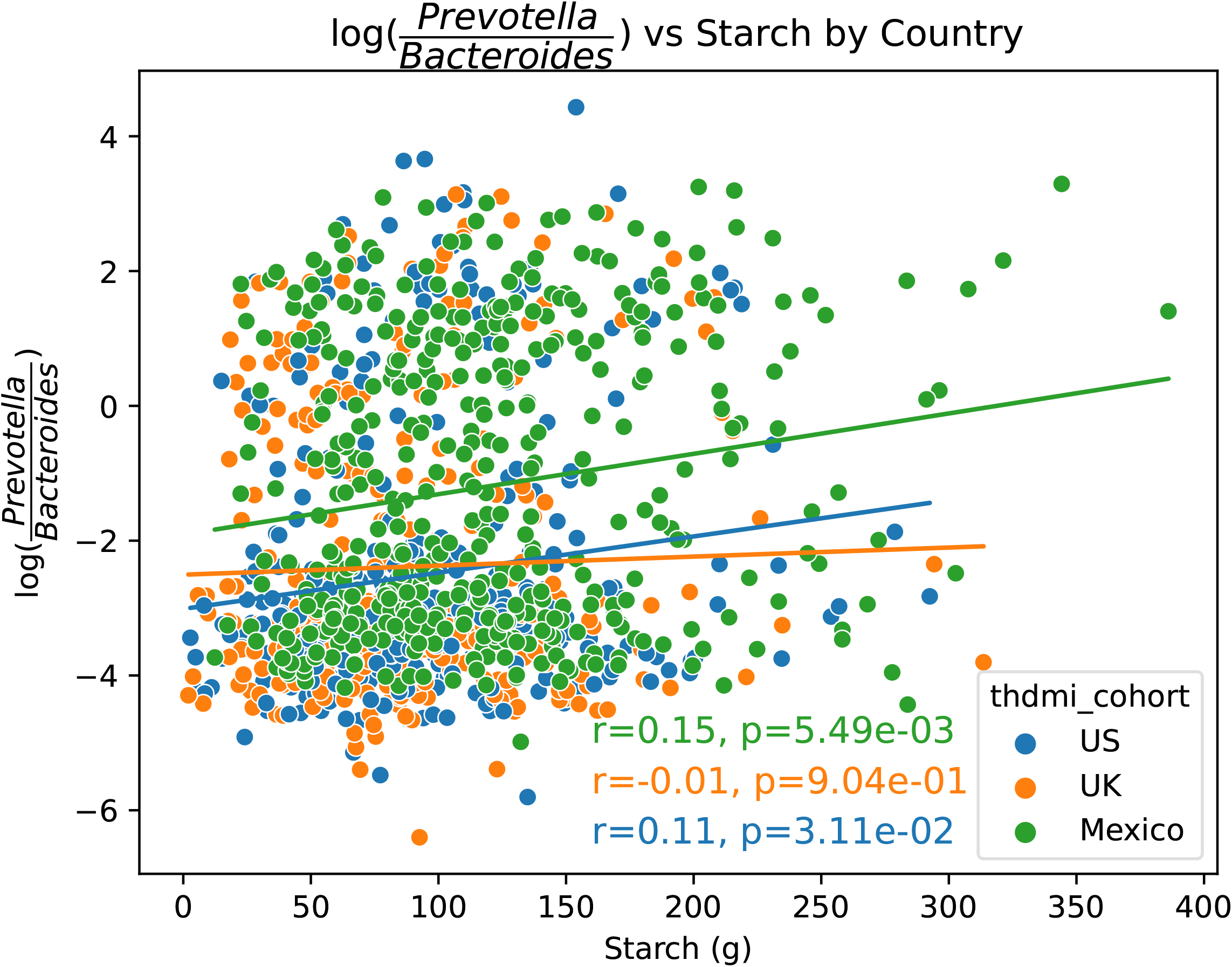
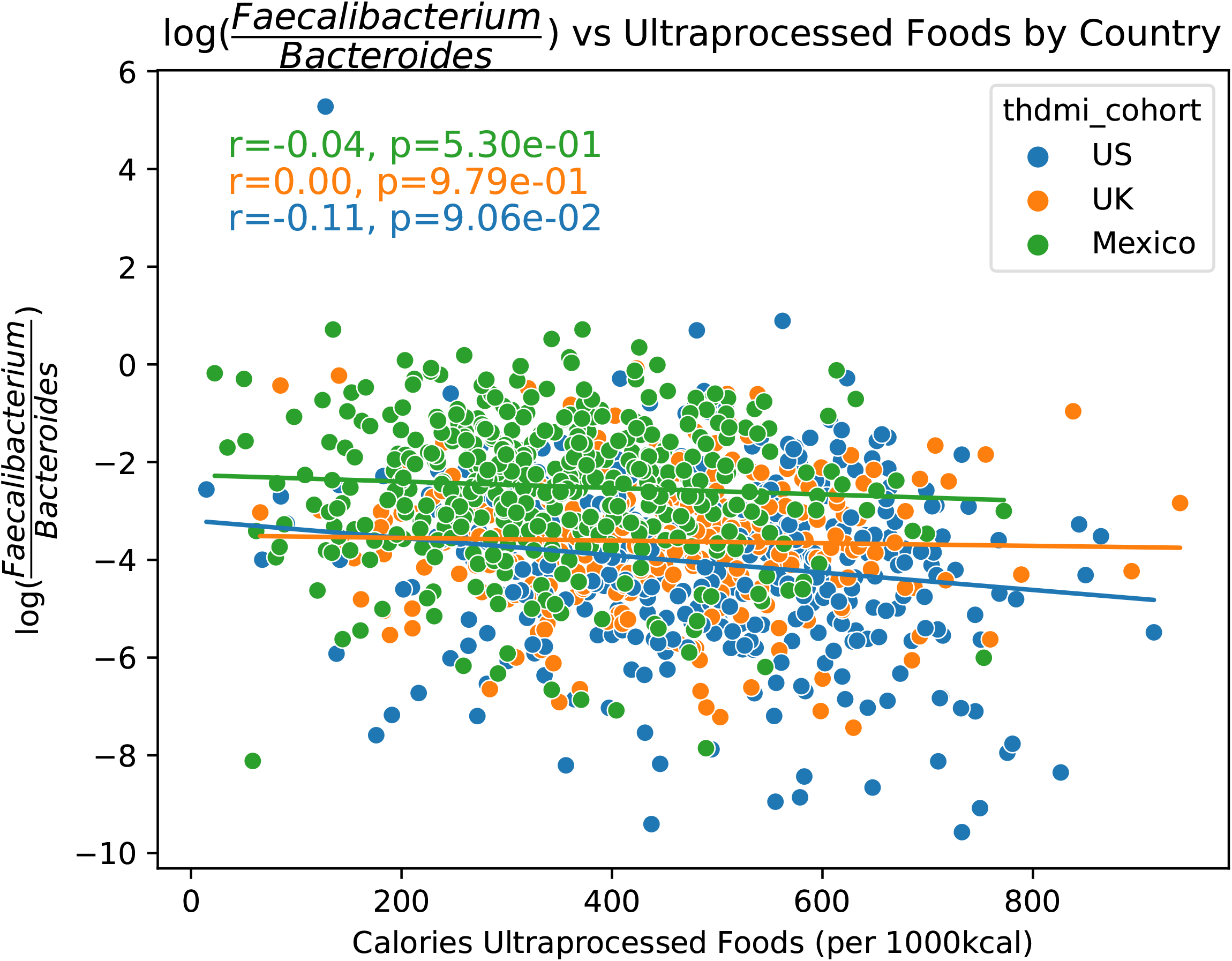
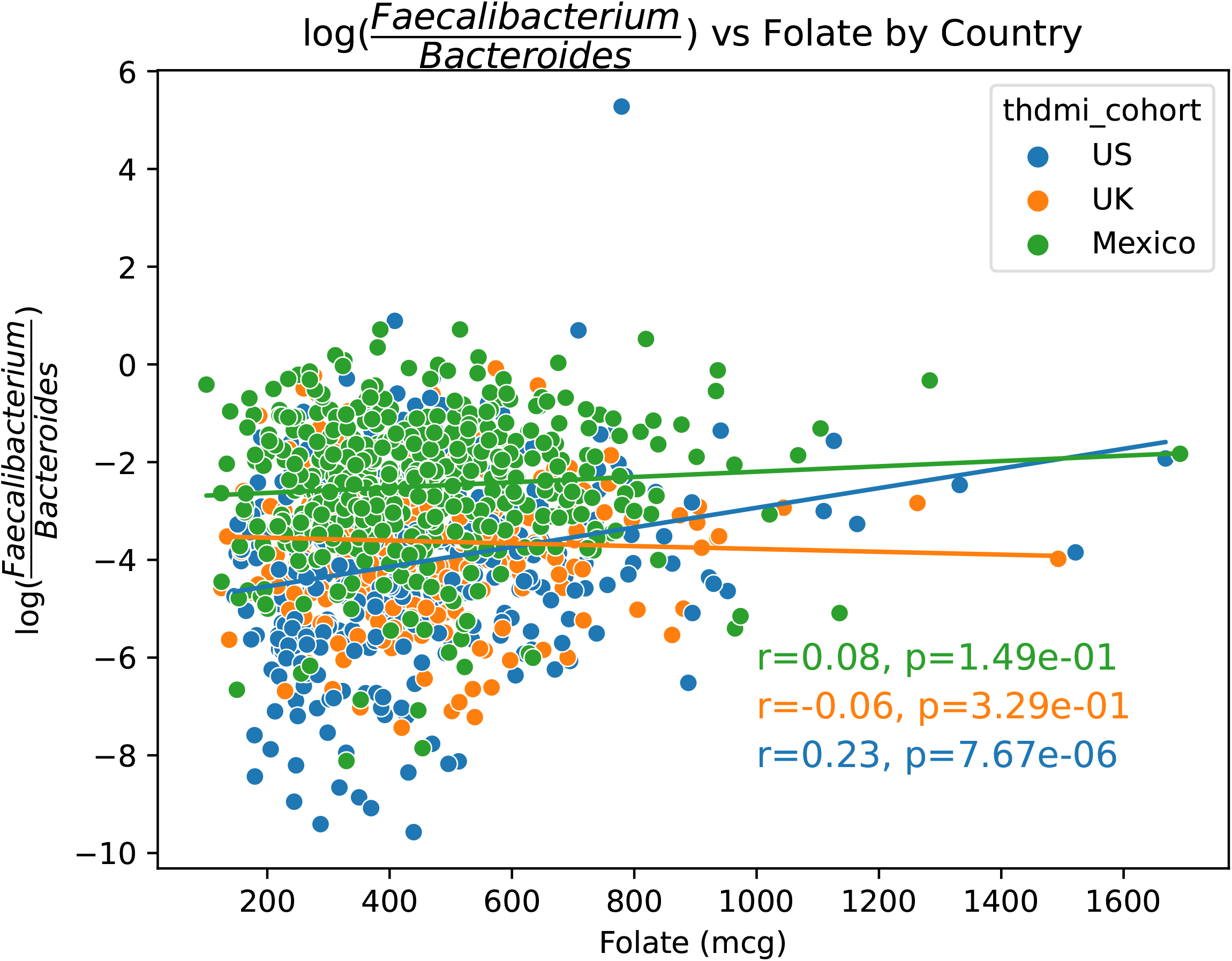
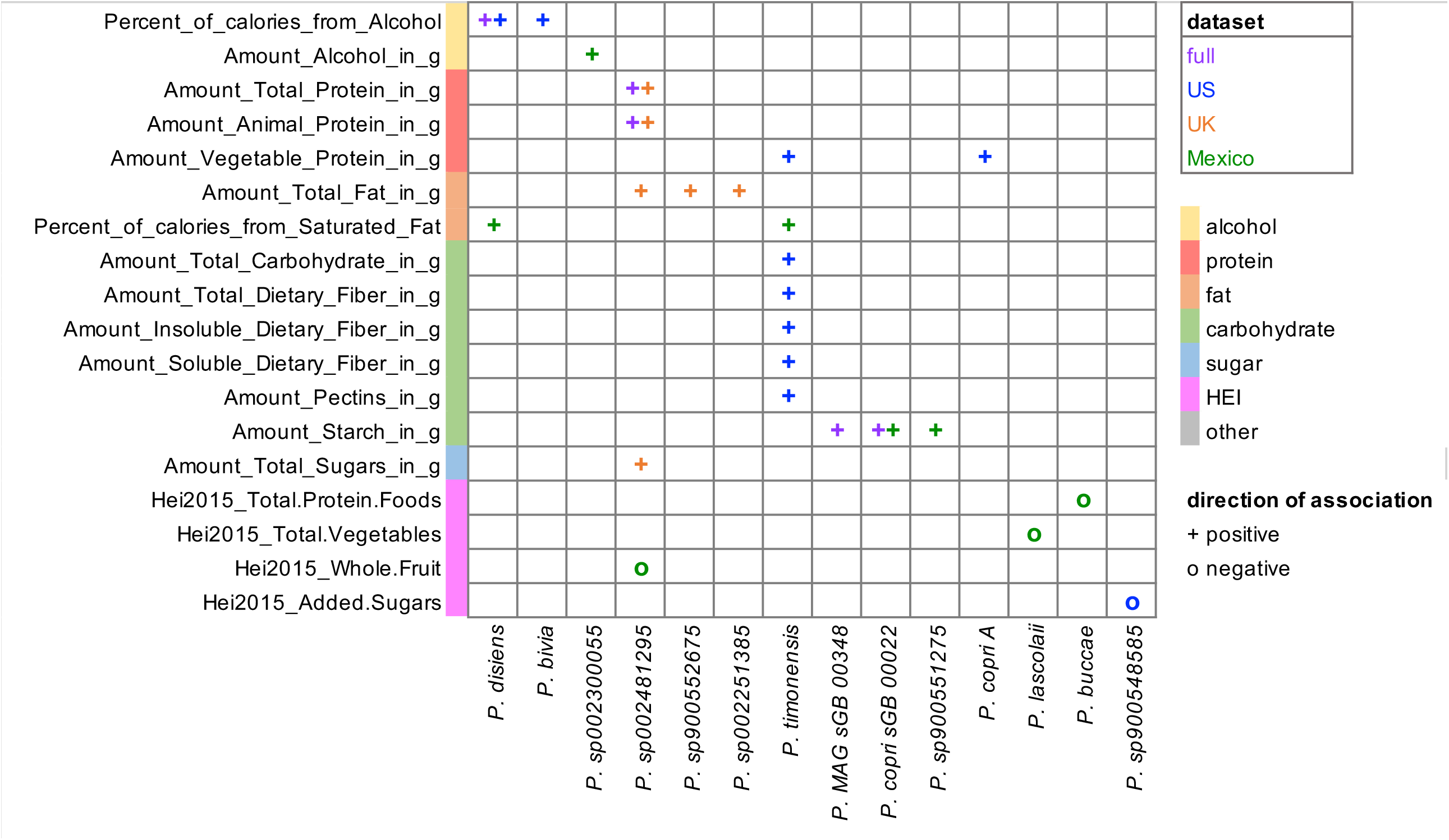
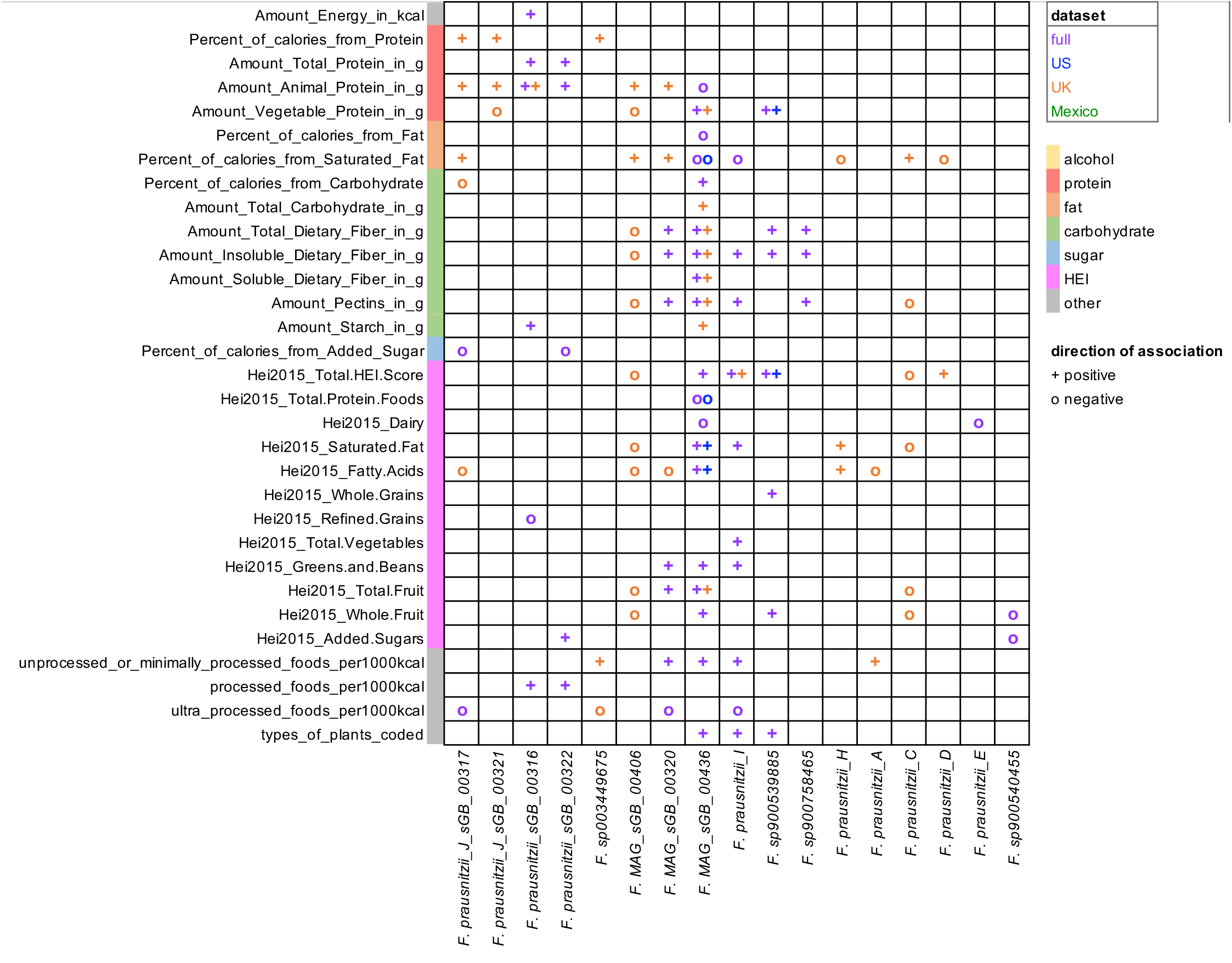

## References

1. McDonald D, Hyde E, Debelius JW, Morton JT, Gonzalez A, Ackermann G, Aksenov AA, Behsaz B, Brennan C, Chen Y. 2018. American gut: an open platform for citizen science microbiome research. Msystems 3:10.1128/msystems.00031-18.

2. Kristal AR, Kolar AS, Fisher JL, Plascak JJ, Stumbo PJ, Weiss R, Paskett ED. 2014. Evaluation of web-based, self-administered, graphical food frequency questionnaire. Journal of the Academy of Nutrition and Dietetics 114:613–621.

3. Cotillard A, Cartier-Meheust A, Litwin NS, Chaumont S, Saccareau M, Lejzerowicz F, Tap J, Koutnikova H, Lopez DG, McDonald D. 2022. A posteriori dietary patterns better explain variations of the gut microbiome than individual markers in the American Gut Project. The American journal of clinical nutrition 115:432–443.

4. Villarroel M BD, & Jen A. 2019. Tables of Summary Health Statistics for U.S. Adults: 2018 National Health Interview Survey. National Center for Health Statistics, http://www.cdc.gov/nchs/nhis/SHS/tables.htm.

5. (CDC) CfDCaP. 2017-2020. National Health and Nutrition Examination Survey (NHANES). Hyattsville, MD. https://www.cdc.gov/nchs/nhanes/index.htm.

6. Armstrong G, Martino C, Morris J, Khaleghi B, Kang J, DeReus J, Zhu Q, Roush D, McDonald D, Gonazlez A. 2022. Swapping metagenomics preprocessing pipeline components offers speed and sensitivity increases. Msystems 7:e01378–21.

7. Zhu Q, Huang S, Gonzalez A, McGrath I, McDonald D, Haiminen N, Armstrong G, Vázquez-Baeza Y, Yu J, Kuczynski J. 2022. Phylogeny-aware analysis of metagenome community ecology based on matched reference genomes while bypassing taxonomy. Msystems 7:e00167–22.

8. Martino C, Morton JT, Marotz CA, Thompson LR, Tripathi A, Knight R, Zengler K. 2019. A novel sparse compositional technique reveals microbial perturbations. MSystems 4:10.1128/msystems.00016-19.

9. Sanders JG, Sprockett DD, Li Y, Mjungu D, Lonsdorf EV, Ndjango J-BN, Georgiev AV, Hart JA, Sanz CM, Morgan DB. 2023. Widespread extinctions of co-diversified primate gut bacterial symbionts from humans. Nature microbiology 8:1039–1050.

10. Olm MR, Brown CT, Brooks B, Banfield JF. 2017. dRep: a tool for fast and accurate genomic comparisons that enables improved genome recovery from metagenomes through de-replication. The ISME journal 11:2864–2868.

11. Olm MR, Crits-Christoph A, Bouma-Gregson K, Firek BA, Morowitz MJ, Banfield JF. 2021. inStrain profiles population microdiversity from metagenomic data and sensitively detects shared microbial strains. Nature Biotechnology 39:727–736.

12. Benus RF, van der Werf TS, Welling GW, Judd PA, Taylor MA, Harmsen HJ, Whelan K. 2010. Association between Faecalibacterium prausnitzii and dietary fibre in colonic fermentation in healthy human subjects. British journal of nutrition 104:693–700.

13. Kovatcheva-Datchary P, Nilsson A, Akrami R, Lee YS, De Vadder F, Arora T, Hallen A, Martens E, Björck I, Bäckhed F. 2015. Dietary fiber-induced improvement in glucose metabolism is associated with increased abundance of Prevotella. Cell metabolism 22:971–982.

14. Prasoodanan PK V, Sharma AK, Mahajan S, Dhakan DB, Maji A, Scaria J, Sharma VK. 2021. Western and non-western gut microbiomes reveal new roles of Prevotella in carbohydrate metabolism and mouth–gut axis. npj Biofilms and Microbiomes 7:77.

15. De Filippis F, Pellegrini N, Laghi L, Gobbetti M, Ercolini D. 2016. Unusual sub-genus associations of faecal Prevotella and Bacteroides with specific dietary patterns. Microbiome 4:1–6.

16. Ahrens AP, Culpepper T, Saldivar B, Anton S, Stoll S, Handberg EM, Xu K, Pepine C, Triplett EW, Aggarwal M. 2021. A six-day, lifestyle-based immersion program mitigates cardiovascular risk factors and induces shifts in gut microbiota, specifically lachnospiraceae, ruminococcaceae, faecalibacterium prausnitzii: a pilot study. Nutrients 13:3459.

17. Nadkarni M, Browne G, Chhour K-L, Byun R, Nguyen K-A, Chapple C, Jacques N, Hunter N. 2012. Pattern of distribution of Prevotella species/phylotypes associated with healthy gingiva and periodontal disease. European journal of clinical microbiology & infectious diseases 31:2989–2999.

18. Alpizar-Rodriguez D, Lesker TR, Gronow A, Gilbert B, Raemy E, Lamacchia C, Gabay C, Finckh A, Strowig T. 2019. Prevotella copri in individuals at risk for rheumatoid arthritis. Annals of the rheumatic diseases 78:590–593.

19. Guevara-Cruz M, Flores-López AG, Aguilar-López M, Sánchez-Tapia M, Medina-Vera I, Díaz D, Tovar AR, Torres N. 2019. Improvement of lipoprotein profile and metabolic endotoxemia by a lifestyle intervention that modifies the gut microbiota in subjects with metabolic syndrome. Journal of the American Heart Association 8:e012401.

20. Martín R, Rios-Covian D, Huillet E, Auger S, Khazaal S, Bermúdez-Humarán LG, Sokol H, Chatel J-M, Langella P. 2023. Faecalibacterium: a bacterial genus with promising human health applications. FEMS Microbiology Reviews 47:fuad039.

21. Blanco-Míguez A, Gálvez EJ, Pasolli E, De Filippis F, Amend L, Huang KD, Manghi P, Lesker T-R, Riedel T, Cova L. 2023. Extension of the Segatella copri complex to 13 species with distinct large extrachromosomal elements and associations with host conditions. Cell host & microbe 31:1804–1819. e9.

22. Sakamoto M, Sakurai N, Tanno H, Iino T, Ohkuma M, Endo A. 2022. Genome-based, phenotypic and chemotaxonomic classification of Faecalibacterium strains: proposal of three novel species Faecalibacterium duncaniae sp. nov., Faecalibacterium hattorii sp. nov. and Faecalibacterium gallinarum sp. nov. International Journal of Systematic and Evolutionary Microbiology 72:005379.

23. De Filippis F, Pasolli E, Ercolini D. 2020. Newly explored Faecalibacterium diversity is connected to age, lifestyle, geography, and disease. Current Biology 30:4932–4943. e4.

